# Chronic Alcohol Exposure Induces Persistent DNA Damage Response Signaling and Sensitizes Breast Epithelial Cells to PARP Inhibition

**DOI:** 10.64898/2026.09.21.753230

**Authors:** Ming Zhao, Zhikun Ma, Amanda B Parris, Jadyn Beacham, Xiaohe Yang

**Affiliations:** Biomedical/Biotechnology Research Institute, Department of Biological and Biomedical Sciences, North Carolina Central University, Kannapolis, NC 28081 USA

**Keywords:** Alcohol, PARP inhibitor, DNA damage, sensitization, breast epithelial cells

## Abstract

Chronic alcohol consumption is associated with increased breast cancer risk; however, the long-term effects of alcohol exposure on DNA damage response (DDR) signaling and therapeutic vulnerability in breast epithelial cells remain incompletely understood. In the present study, we established a chronic alcohol exposure model using immortalized human breast epithelial MCF-10A cells to investigate persistent alterations in DNA damage signaling and sensitivity to poly (ADP-ribose) polymerase (PARP) inhibition. MCF-10A cells that chronically exposed to 0.2% alcohol for 20 weeks and then withdrew alcohol treatment for 4 weeks still retained elevated γH2AX levels and sustained activation of ATM/ATR-associated DDR pathways. Chronic alcohol-exposed cells also exhibited increased sensitivity to the PARP inhibitors olaparib and veliparib, as demonstrated by reduced cell viability, decreased clonogenic survival, and enhanced apoptosis. Mechanistically, olaparib induced greater G2/M checkpoint activation and replication-associated DNA damage in chronic alcohol-exposed cells compared with control cells. In addition, chronic alcohol exposure increased PARP expression and enhanced activation of DDR signaling following PARP inhibition. These findings demonstrate that chronic alcohol exposure induces persistent DNA repair stress and sensitizes breast epithelial cells to PARP inhibition. Our study suggests that chronic alcohol-associated genomic stress may create potential vulnerability to PARP inhibition.

**Highlights:**

- Chronic alcohol exposure induces persistent DNA damage response signaling in MCF-10A cells.
- Chronic alcohol-exposed cells exhibit enhanced sensitivity to the PARP inhibitors.
- Chronic alcohol exposure increased G2/M arrest induced by PARP inhibitor and replication-associated DNA damage.
- Chronic alcohol exposure upregulates PARP expression and alters DDR signaling responses.

## Introduction

Alcohol consumption is a well-established risk factor for breast cancer and has been associated with increased incidence and progression of multiple malignancies [1]. Epidemiological studies have shown that even moderate alcohol intake is associated with elevated breast cancer risk, although the molecular mechanisms underlying alcohol-mediated carcinogenesis remain incompletely understood [2,3]. Ethanol metabolism generates acetaldehyde and reactive oxygen species, both of which can induce DNA damage and genomic instability. Accumulating evidence indicates that alcohol exposure promotes DNA adduct formation, replication stress, chromosomal rearrangements, and defects in genome maintenance pathways [4,5]. DNA damage response (DDR) pathways play critical roles in maintaining genomic integrity following replication-associated stress and DNA lesions. Upon DNA damage, signaling networks involving ATM, ATR, Chk1, Chk2, and p53 coordinate cell cycle checkpoint activation and DNA repair processes [6]. Chronic activation or dysregulation of these pathways is increasingly recognized as a hallmark of early tumorigenic transformation and therapeutic vulnerability [7]. Previous studies have demonstrated that acetaldehyde-induced DNA lesions interfere with DNA replication fork progression and stimulate homologous recombination-associated repair responses [8]. However, whether chronic alcohol exposure induces persistent alterations in DDR signaling in breast epithelial cells remains poorly understood.

Poly (ADP-ribose) polymerase (PARP) enzymes are central regulators of DNA repair and replication-associated damage responses. Pharmacological inhibition of PARP has emerged as an effective therapeutic strategy in tumors with homologous recombination deficiencies, particularly BRCA1/2-mutated cancers [9]. In addition to canonical BRCA-deficient contexts, increasing evidence suggests that chronic replication stress and persistent DNA repair abnormalities may also enhance cellular sensitivity to PARP inhibition [10]. Nevertheless, the impact of chronic alcohol exposure on PARP inhibitor responsiveness has not been clearly defined.

In the present study, we established a chronic alcohol exposure model using immortalized human breast epithelial MCF-10A cells to investigate the long-term effects of alcohol on DNA damage signaling and therapeutic vulnerability. We demonstrate that chronic alcohol exposure induces persistent activation of DDR pathways, enhances replication-associated DNA damage, and increases sensitivity to the PARP inhibitors olaparib and veliparib. Our findings suggest that chronic alcohol exposure generates sustained DNA repair stress that may create a therapeutically exploitable vulnerability to PARP inhibition.

## Methods and Materials

### 1. Cell culture

MCF-10A cells were ordered from American Type Culture Collection (ATCC, Manassas, USA). The cells were cultured in DMEM/F-12 medium supplemented with 10% fetal bovine serum (FBS), penicillin (100 U/mL), and streptomycin (100 μg/mL), and plus cholera toxin (100 ng/ml), hydrocortisone (1 ug/ml), Insulin (10 ug/ml), EGF (10 ng/ml). For the chronic alcohol exposure (the cells were named by MCF-10A/CAE), the cells were treated with 0.2% alcohol (200 proof) for 20 weeks and withdrew alcohol treatment for 4 weeks. The control cells, which were named by MCF-10A/C, were passaged with the same procedures without alcohol treatment.

### 2. Cell viability assay

Cell viability was measured using Cell Counting Kit-8 (CCK-8; Enzo, Farmingdale, USA). The cells were seeded into 96-well plates (1 × 103 cells per well) for 24 h prior to treatment. The cells were then treated with indicated agents at different concentrations for 96 h. At the endpoint, CCK-8 solution was added to the wells (10 μl per well) and incubated for 2 h. The cell survival fractions were calculated based on absorbance (A) at 450 nm measured with a microplate reader (SynergyMx, BioTek), and the data was processed using GraphPad Prism 8 software.

### 3. Colony formation assay

Cells were seeded into 6-well plates at a density of 800 cells per well. At the second day, the cells were treated with indicated reagent at different concentrations for ten days. The medium was replaced every three days during the treatment. At the endpoint, the cells were washed with PBS twice and fixed with acetone/methanol (1:1) for 5 minutes. After staining with 0.5% crystal violet for 20 minutes, the plates were washed with deionized water and air-dried. Images were captured with a Nikon stereoscopic microscope (SMZ-745T) and analyzed with ImageJ software.

### 4. Apoptosis ELISA

Cell apoptosis was measured by Cell Death Detection ELISA kit (Roche Life Science, Indianapolis, USA), in which the cell death quantitatively determined by cytoplasmic histone-associated DNA fragments (mono- and oligonucleosomes). After treatment, the cells were collected by trypsinization and counted. Ten thousand cells in each sample were incubated with 200 μl lysing buffer for 30 min and centrifugated at 200 × *g* for 10 min. Then, 20 μl of supernatant was transferred into a microplate for the reaction with the immunoreagents. The washed wells were incubated with ABTS substrate solution and ABTS stop solution for color development, and then followed by absorbance reading at 405 nm using a microplate reader. Relative apoptosis based on triplicate samples was presented as a ratio of the apoptosis in drug treated groups over the control group.

### 5. Live/Dead cell counting by ImageXpress Pico

The cells were plated in a 96-well black, clear-bottom microplates at 2000 cells per well. At the second day, the cells were treated with indicated doses of Olaparib for 48 h. Then, the cells with EarlyTox Live/Dead assay kit (Molecular Devices, R8341). The cells were incubated with Calcein AM and EthD-Ⅲ (working concentrations were both 1 μM), and the plates were incubated at 37°C, 5% CO2 for 45 minutes. Immediately after the final incubation, the plates were imaged on the ImageXpress Pico system using a 4X Plan Apo objective and the FITC and Texas Red filter sets. The cell counting was analyzed by the Automated Cell Imaging System.

### 6. Cell cycle analysis

The cells were collected by trypsinization and washed with PBS. The cells were fixed drop-wise with 70% ethanol and stored at −20°C overnight. The samples were then washed with ice-cold PBS twice, followed by incubation with the staining buffer (PI 33 μg/mL, 0.1% Triton X-100, 500 μg/mL RNase A) at 37 °C for 30 min. The cells were then analyzed using a Guava EasyCyte 8 flow cytometer (Millipore, MA, USA). The DNA content and cell cycle phases of individual samples were analyzed with ModFit LT Software

### 7. Immunofluorescence (IF)

The cells were fixed in 4% paraformaldehyde for 15 min at room temperature. After washing with PBS, the cells were permeabilized with 0.1% Triton X-100 for 20 min and continually blocked with 3% BSA (in PBS) for 30 min. Cells were then incubated with primary antibodies diluted in 3% BSA (PBS) overnight at 4 °C. After washing three times, Alexa Fluor 488/546-labeled secondary antibodies (Thermo Fisher Scientific) diluted in PBS were added and incubated for 1 h at room temperature in the dark. After washing three times, the cells were mounted in Antifade Mounting Medium containing DAPI (VecorLabs). Images were acquired using a Nikon fluorescence microscope. For CIdU labeling, living cells were treated with CIdU (25 μM) for 20 min at 37°C in a 5% CO2 incubator, and then followed by IF staining to assess DNA damage in cells undergoing DNA replication. The primary antibody of Rat monoclonal anti-BrdU antibody [BU1/75 (ICR1)] (Abcam, ab6326) and the corresponding secondary antibody of Alexa Fluor 555 goat anti-rat IgG (Thermo Fisher Scientific, A21434) were applied.

### 8. Quantitative real time-PCR (*q*-RT-PCR)

Total RNA was extracted using TRIzol (Invitrogen, Carlsbad, CA) according to the manufacturer’s protocol and quantified using a NanoDrop1000 spectrophotometer (Thermo Fisher). First-stand cDNA of each sample was synthesized using an All-in-one First-Strand cDNA Synthesis Kit (Bio-Rad). Quantitative real-time PCR was performed on a PCR cycler (Bio-Rad CFX96) with synthetic primers (IDT, Coralville, Iowa). The primers for PARP are F: 5’-tggaacatcaaggacgagct-3’ and R: 5’-catcgctcttgaagaccagc-3’; The primers for ACTB are F: 5’-catccgcaaagacctgtacg-3’ and R: 5’-cctgcttgctgatccacatc-3’. Samples were subjected to the following reaction conditions: 95 °C for 3 min, followed by 45 cycles of 95 °C for 10 sand 55 °C for 30 s. The 2^-ΔΔCt^ method was used to calculate relative mRNA levels. The expression of ACTB served as an internal control.

### 9. Western blot

Total proteins were extracted with NP-40 buffer (50 mM Tris-HCl pH=7.6, 150 mM NaCl, 1% NP-40, 5 mM EDTA) supplemented protease and phosphatase inhibitor cocktails (Thermo Scientific, IL, USA) and centrifuged at 12000 rpm for 15 min at 4 °C. BCA protein assay kit (Thermo Scientific, IL, USA) was used to determine the protein concentration of the supernatant. Equal amounts of 20 μg of protein were separated by sodium dodecyl sulfate-polyacrylamide gel electrophoresis (SDS-PAGE) and electrically transferred onto a polyvinylidene difluoride (PVDF) membrane (Millipore, MA, USA). After nonspecific binding was blocked with 5% nonfat milk in Tris-buffered saline/Tween 20 (TBST) at room temperature for 2 h, membranes were incubated with primary antibodies. The primary antibodies, PARP (#9542), p-p53(#9284), p-H2AX (#9718), p-ATM(#5883), ATM (#2873), p-ATR (#2853), ATR (#2790), p-Chk1 (#2348), Chk1 (#2360), p-Chk2 (#2197), Chk2 (#6334), Ku80 (#2180), Rad50 (#3427), p-Cdc2 (#4539), Cdc2 (#9116), Cdc25C (#4688), Cyclin B (#12231), BRCA1 (#9010), BRCA2 (#26542) and HRP-linked secondary antibodies were ordered from Cell Signaling Technology (Danvers, MA). The primary antibodies, p53 (sc-126), E2F1 (sc-251) and β-Actin (sc-47778) were ordered from Santa Cruz Biotechnology (Dallas, TX). After three washes with TBST, membranes were incubated with HRP-conjugated secondary antibody diluted at 1:2000 at room temperature for 2 h. After washing three times, these protein blots were detected on Azure biosystems.

### 10. Statistical analysis

The quantitative results were analyzed with GraphPad Prism software. The data are presented as means ± S.E. Two-group comparisons were performed using Student’s t-test, while one-way ANOVA was used for multiple group comparisons. Statistical significance was defined as * p < 0.05, ** p < 0.01.

## Results

### 1. Chronic alcohol exposure induces persistent DNA damage response in MCF-10A cells

Prior to investigating the effects of chronic alcohol exposure on DNA damage in non-transformed breast epithelial MCF-10A cells, we first examined the response to acute alcohol treatment. MCF-10A cells were exposed to increasing concentrations of alcohol (0.1–0.4%) for 2 h and analyzed for DNA damage-associated responses. Immunofluorescence analysis of γH2AX foci, a marker of DNA damage, revealed a dose-dependent increase following alcohol treatment (Fig. 1A, B). Consistent with these findings, Western blot analysis showed elevated γH2AX levels and increased phosphorylation of p53, whereas total p53 protein levels increased only modestly (Fig. 1C). Together, these results indicate that acute alcohol exposure induces DNA damage and activates the DNA damage response (DDR) pathway in MCF-10A cells.

**Figure 1.**
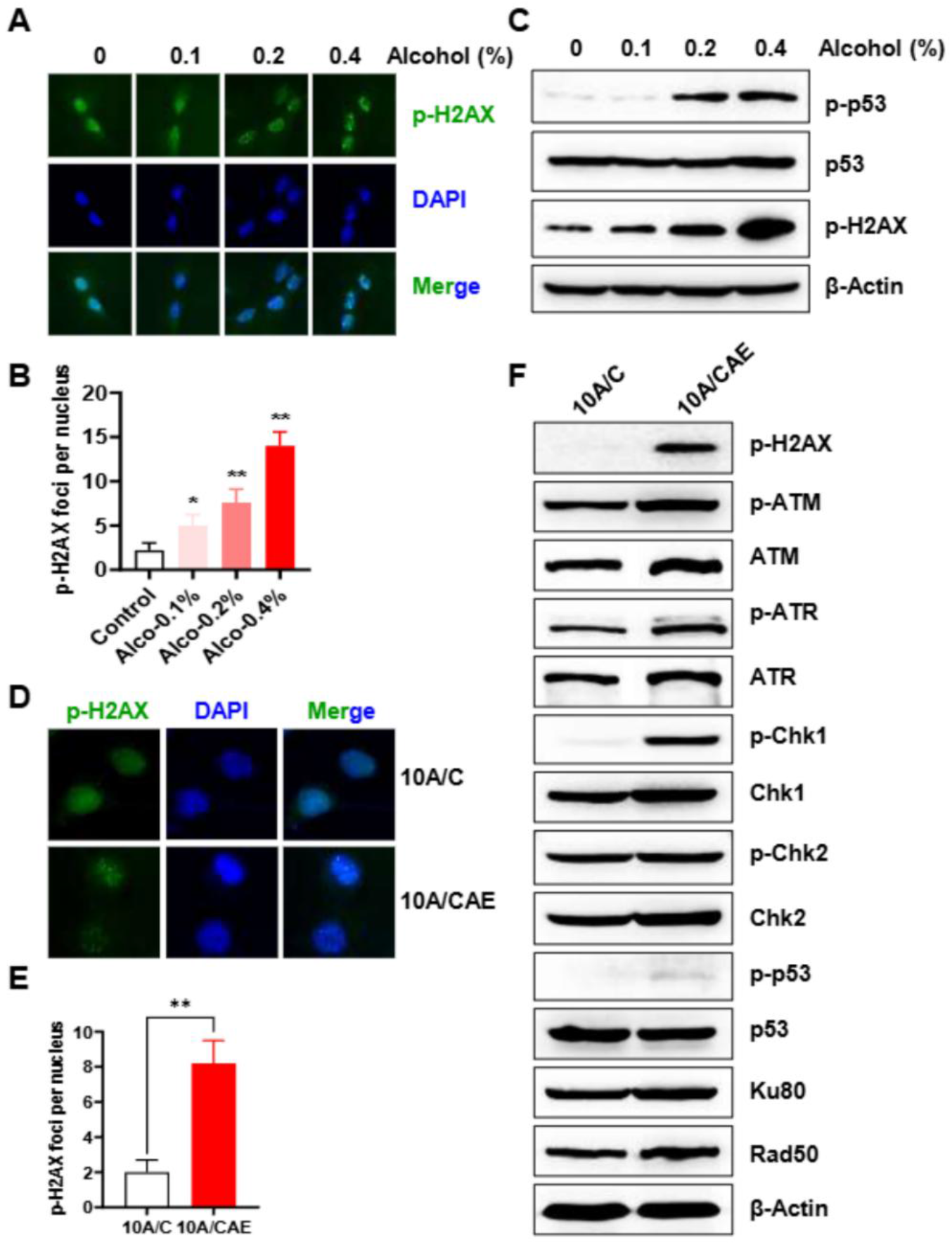
Acute alcohol exposure induces DNA damage in MCF-10A cells. **A**. MCF-10A cells were treated with different concentrations of alcohol (0, 0.1%, 0.2%, 0.4%) for 2 h. Then, the cells were fixed and stained with anti-p-H2AX antibody, and then subjected to immunofluorescent microscopy. The representative images from three independent experiments were shown. **B**. p-H2AX foci were quantified as foci number per nucleus in each group. **p < 0.01. **C**. MCF-10A cells were treated as above described. The protein levels of p-p53, p53, p-H2AX were detected by western blotting. β-Actin serves as loading control. **D**. The representative images of MCF-10A control (10A/C) and chronic 0.2% alcohol-exposed 20 weeks (10A/CAE) cells underwent immunofluorescent staining with anti-p-H2AX antibody. **E**. The bar graph shows the quantitation of p-H2AX foci number per nucleus in each group. **F**. Protein lysates were prepared from both control (10A/C) and chronic alcohol exposed (10A/CAE) MCF-10A cells, followed by western blotting. Protein levels of indicated markers were analyzed. β-Actin serves as loading control.

To investigate the effects of chronic alcohol exposure on persistent DNA damage for subsequent functional analyses, MCF-10A cells were continuously cultured in the presence of 0.2% alcohol for 20 weeks, followed by maintenance in alcohol-free medium for at least 4 weeks before analysis. The resulting cells were designated MCF-10A/CAE, while passage-matched untreated cells served as controls (MCF-10A/C). Under basal alcohol-free culture conditions, MCF-10A/CAE cells exhibited persistent γH2AX foci formation compared with control cells (Fig. 1D, E), suggesting sustained DNA damage or unresolved DNA lesions following chronic alcohol exposure. Western blot analysis further revealed increased phosphorylation of ATM, ATR, Chk1, Chk2, and p53 in MCF-10A/CAE cells (Fig. 1F). Expression of the DNA repair-associated proteins Ku80 and Rad50 was also elevated. Together, these findings indicate that chronic alcohol exposure induces persistent activation of DNA damage response (DDR) signaling pathways in MCF-10A cells.

### 2. MCF-10A/CAE cells exhibit increased sensitivity to PARP inhibitors

Emerging evidence suggests that DNA repair defects beyond BRCA1/2 mutations can confer sensitivity to PARP inhibitors. Given that chronic alcohol exposure induced persistent DNA damage and alterations in DNA repair-associated proteins in MCF-10A/CAE cells, we examined their response to the PARP inhibitors olaparib and veliparib in the absence of alcohol. Cell viability assays showed that MCF-10A/CAE cells were more sensitive to both olaparib and veliparib than control cells, particularly at higher drug concentrations (Fig. 2A, B). Consistently, clonogenic assays demonstrated significantly reduced colony-forming ability in olaparib-treated MCF-10A/CAE cells compared with control cells (Fig. 2C, D). To further evaluate cell death responses, paired cell lines were treated with olaparib and analyzed using the ImageXpress Pico high-content imaging system. Olaparib induced substantially greater cell death in MCF-10A/CAE cells than in control cells (Fig. 2E-G). Consistent with these findings, apoptosis ELISA analysis revealed significantly higher levels of apoptosis in olaparib-treated MCF-10A/CAE cells (Fig. 2H). Together, these results demonstrate that chronic alcohol exposure increases sensitivity to PARP inhibition, suggesting persistent DDR activation der MCF-10A/CAE cells more dependent on PARP-mediated DNA repair pathways.

**Figure 2.**
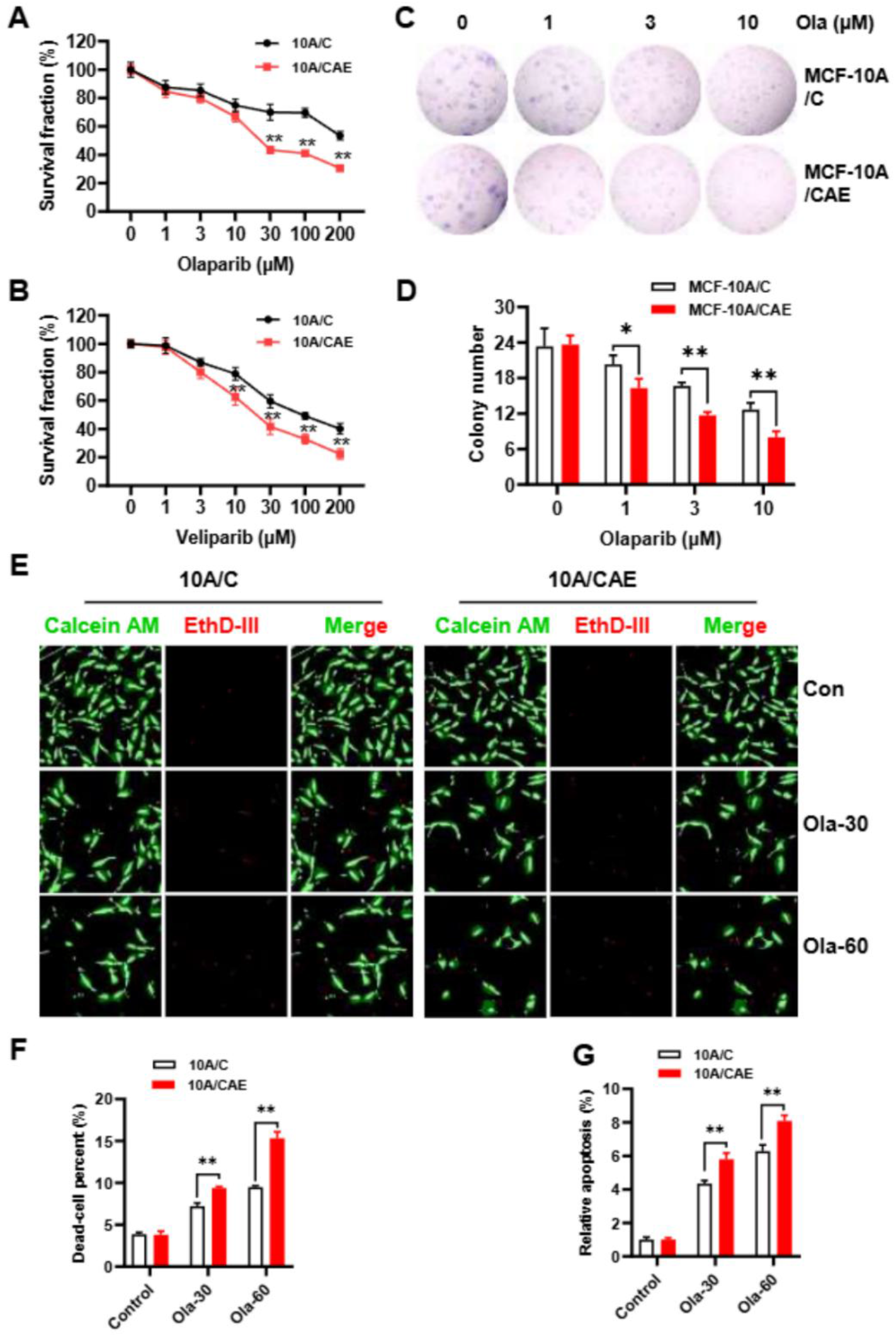
Chronic alcohol exposure renders MCF-10A cells sensitive to PARP inhibitors. **A&B**. MCF-10A/C and MCF-10A/CAE cells were treated with different doses of olaparib or veliparib (0, 1, 3, 10, 30, 100, 200 μM) for 4 days, followed by CCK-8 assay. The values were expressed as means ± SEM of three independent replicates. **p<0.01 compared with control cells treated with the same dose of inhibitor. **C&D**. Clonogenic assays of MCF-10A/C and MCF-10A/CAE cells. The paired cell lines were seeded in 6-well plates and treated with olaparib (0, 1, 3, 10 μM) for 10 days. The colonies were stained with crystal violet. The representative images were shown (B). The bar graph shows colony number in each group of triple replicates. *p<0.05 **p<0.01. E-G. Live/Dead cell counting by ImageXpress Pico. MCF-10A/C and MCF-10A/CAE cells were treated with olaparib (0, 30, 60 μM) for 3 days, and followed Live/Dead cell counting by ImageXpress Pico. Representative images of live cells stained with calcein AM (green) and dead cells stained with EthD-Ⅲ (Red) were shown **(E)**. The percent of live cells **(F)** and dead cells **(G)** were quantified respectively. *p<0.05 **p<0.01. **H**. MCF-10A/C and MCF-10A/CAE cells were treated with olaparib (0, 30, 60 μM) for 3 days. Then, cell apoptosis in each group was determined using apoptosis ELISA kit. The values were expressed as means ± SEM of three independent replicates. **p<0.01.

### 3. Olaparib induces enhanced G2/M arrest and replication-associated DNA damage in chronic alcohol-exposed cells

To investigate the mechanisms underlying increased PARP inhibitor sensitivity, cell cycle distribution was examined following olaparib treatment. Under basal conditions, MCF-10A/CAE cells displayed a modest increase in S-phase population compared with control cells. Following olaparib treatment, a substantially greater accumulation of cells in the G2/M phase was observed in MCF-10A/CAE cells relative to MCF-10A/C cells (Fig. 3A&B), indicating enhanced checkpoint activation. Consistent with these findings, western blot analysis demonstrated increased phosphorylation of Cdc2, Chk1, and Chk2 in olaparib-treated MCF-10A/CAE cells compared with control cells (Fig. 3C). In contrast, total levels of Cdc2, Cyclin B, and Cdc25C showed relatively limited changes. These data suggest that chronic alcohol exposure enhances olaparib-induced activation of G2/M checkpoint signaling. To determine whether olaparib-induced DNA damage preferentially occurred in replicating cells, cells were pulse-labeled with CIdU prior to immunofluorescence analysis. The p-H2AX-positive nuclei were predominantly detected in CIdU-positive cells following olaparib treatment (Fig. 3D), suggesting that replication-associated DNA damage contributes to the enhanced sensitivity of chronic alcohol-exposed cells to PARP inhibition.

**Figure 3.**
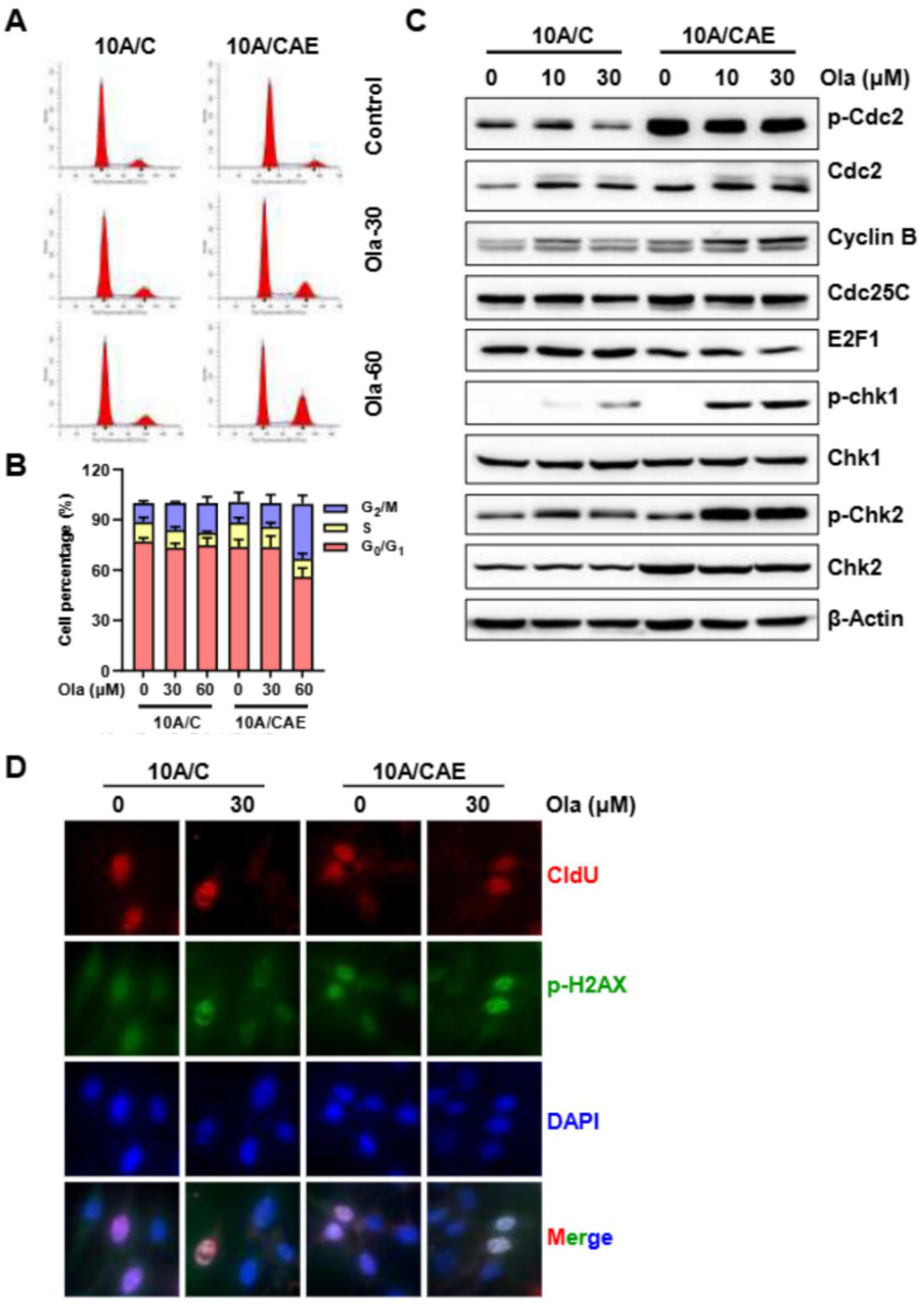
Olaparib induces enhanced G2/M checkpoint activation and replication-associated DNA damage in MCF-10A/CAE cells. **A&B**. MCF-10A/C and MCF-10A/CAE cells were treated with olaparib (0, 30, 60 μM) for 3 days, and followed cell cycle analysis. Representative cell cycle distribution in each group was shown (A). The quantitative cell cycle distribution data was analyzed with ModFit software (B). Data from triplicate samples were reported. **C**. MCF-10A/C and MCF-10A/CAE cells were treated with olaparib (0, 30, 60 μM) for 3 days, and the protein levels of p-Cdc2, Cdc2, Cyclin B and Cdc25c, E2F1, p-Chk1, Chk1, p-Chk2, Chk2, which were involved in cell cycle progression were detected by western blot. β-Actin serves as loading control. **D**. MCF-10A/C and MCF-10A/CAE cells were treated with 30 μM olaparib for 3 days, and then were chased with CIdU (25 μM) for 20 min. The cells were immunofluorescent stained with anti-BrdU antibody (BU1/75 (ICR1)) and anti-H2AX antibody. The representative images were shown.

### 4. Chronic alcohol exposure alters PARP expression and DDR signaling responses

Because PARP is the molecular target of olaparib, we next examined whether chronic alcohol exposure altered PARP expression. Quantitative PCR analysis demonstrated significantly increased PARP mRNA expression in MCF-10A/CAE cells compared with control cells (Fig. 4A). Consistently, western blot analysis confirmed elevated PARP protein expression in chronic alcohol-exposed cells (Fig. 4B). We further examined DDR signaling following olaparib treatment. Although total ATM and ATR protein levels were relatively unchanged, phosphorylation of ATM and ATR was increased to a greater extent in MCF-10A/CAE cells following olaparib exposure (Fig. 4C). Increased expression of BRCA1 and BRCA2 was also observed in alcohol-exposed cells. In addition, phosphorylation of p53 was markedly enhanced in MCF-10A/CAE cells after olaparib treatment, whereas total p53 levels remained relatively stable. These findings indicate that chronic alcohol exposure alters DDR signaling networks and enhances cellular responses to PARP inhibition.

**Figure 4.**
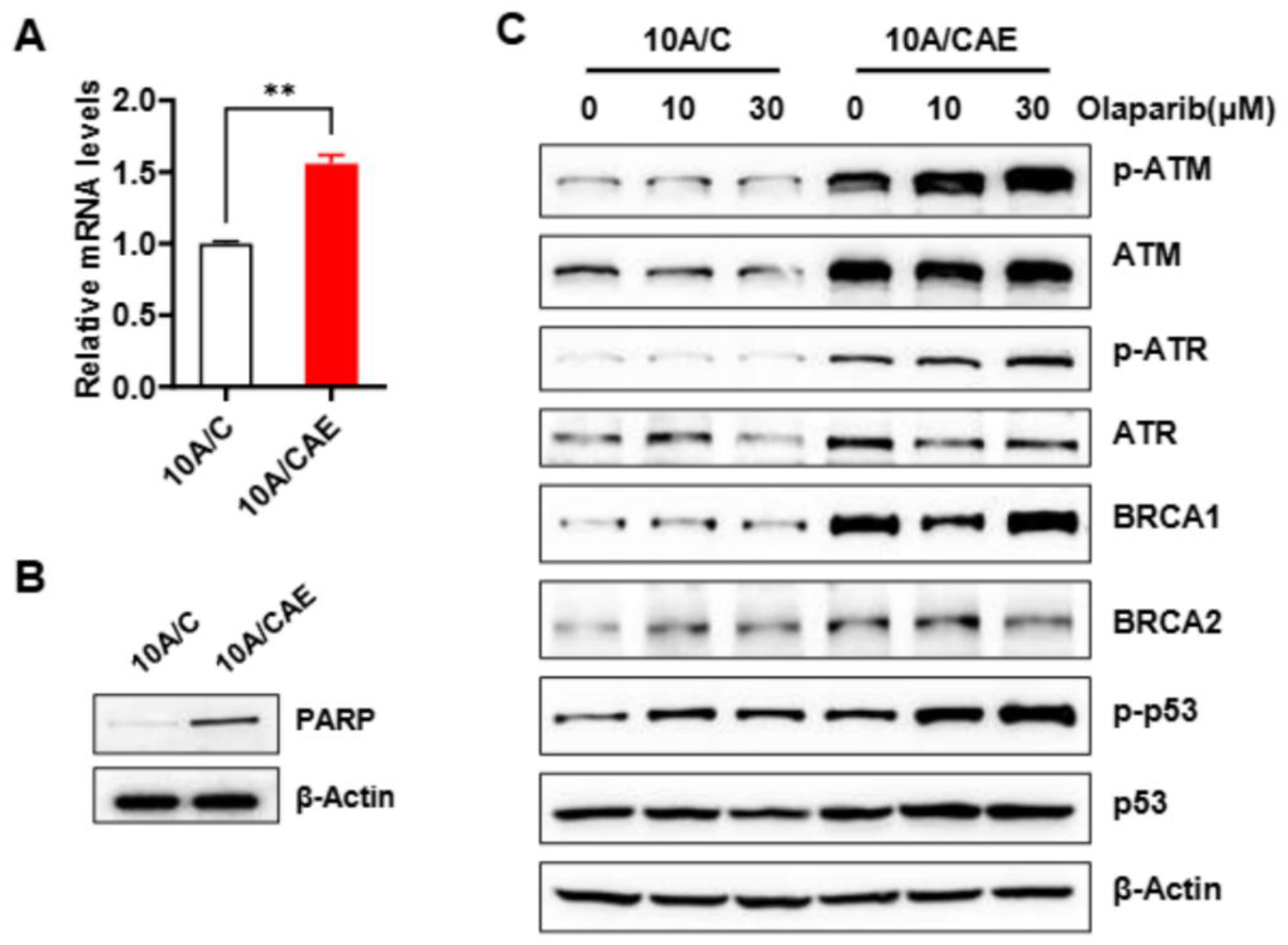
Chronic alcohol exposure altered PARP expression and DDR pathway. **A**. The mRNA levels of PARP in MCF-10A/C and MCF-10A/CAE cells were determined by *q*-PCR assay. **p<0.01. **B**. The protein levels of PARP in MCF-10A/C and MCF-10A/CAE cells were detected with Western blotting. **C**. MCF-10A/C and MCF-10A/CAE cells were treated with olaparib (0, 10, 30 μM) for 3 days. The proteins involved in DDR pathway were detected with western blot.

## Discussion

In the present study, we demonstrate that chronic alcohol exposure induces persistent activation of DNA damage response pathways and enhances sensitivity to PARP inhibition in MCF-10A breast epithelial cells. Using a long-term alcohol exposure model, we found that alcohol-treated cells retained elevated γH2AX (p-H2AX) signaling and sustained activation of ATM/ATR-associated checkpoint pathways even after withdrawal of alcohol exposure. In parallel, chronic alcohol exposure increased cellular sensitivity to the PARP inhibitors olaparib and veliparib, accompanied by enhanced G2/M checkpoint activation, replication-associated DNA damage, and apoptosis. These findings support the concept that chronic alcohol exposure produces persistent DNA repair stress that creates a vulnerability to PARP inhibition.

A notable aspect of this study is the use of non-transformed breast epithelial cells to model chronic alcohol-associated genomic stress. Although alcohol-induced DNA damage has been extensively investigated in cancer cells and in hematopoietic systems, less is known about the long-term effects of chronic alcohol exposure on DNA damage signaling in mammary epithelial cells prior to malignant transformation [11-13]. Our data show that chronic low-dose alcohol exposure induces persistent DDR activation even after prolonged removal of alcohol from the culture conditions, suggesting that alcohol exposure may produce durable alterations in genome maintenance pathways. The persistent elevation of γH2AX, together with increased phosphorylation of ATM, ATR, Chk1, Chk2, and p53, indicates sustained checkpoint signaling and unresolved DNA lesions in chronic alcohol-exposed cells.

Previous studies have demonstrated that acetaldehyde, a major metabolite of ethanol, induces replication fork abnormalities, DNA adduct formation, chromosome rearrangements, and homologous recombination-associated repair responses [14]. Consistent with these observations, our results suggest that replication-associated stress is an important component of the chronic alcohol-induced phenotype. Olaparib-induced γH2AX formation was predominantly observed in CIdU-positive cells, indicating that replicating cells are particularly vulnerable to DNA damage following PARP inhibition. Moreover, chronic alcohol-exposed cells displayed enhanced G2/M accumulation and increased activation of Chk1/Chk2- Cdc2 checkpoint signaling after olaparib treatment. These findings support a model in which chronic alcohol exposure generates persistent replication stress that sensitizes cells to further disruption of DNA repair pathways.

An additional finding of this study is the increased expression of PARP in chronic alcohol-exposed cells. PARP enzymes play critical roles in sensing DNA strand breaks and coordinating repair during replication-associated stress [15]. Elevated PARP expression in MCF-10A/CAE cells may reflect an adaptive response to sustained DNA damage and checkpoint activation. Importantly, despite this apparent compensatory response, chronic alcohol-exposed cells remained more sensitive to pharmacological PARP inhibition. These observations are consistent with the idea that cells experiencing chronic DNA repair stress may become increasingly dependent on PARP-mediated repair pathways for survival [16]. The enhanced sensitivity of chronic alcohol-exposed cells to PARP inhibitors may have broader implications for alcohol-associated carcinogenesis and therapeutic response. PARP inhibitors are primarily used in tumors with homologous recombination deficiencies, particularly BRCA1/2-mutated cancers [17]. However, accumulating evidence suggests that replication stress and persistent DNA repair abnormalities can also confer vulnerability to PARP inhibition in BRCA-proficient settings [18]. Our findings raise the possibility that chronic alcohol exposure may contribute to the development of a PARP inhibitor-sensitive phenotype through sustained DDR activation and replication-associated genomic stress.

We acknowledge that the experiments in this report were performed in a single immortalized breast epithelial cell model, and further validation in additional breast epithelial or breast cancer models will be important. Although chronic alcohol exposure induced persistent DDR activation, the precise molecular mechanisms responsible for increased PARP inhibitor sensitivity remain to be fully defined. Future studies examining replication fork dynamics, homologous recombination activity, and acetaldehyde-mediated DNA lesions may provide additional mechanistic insight. In addition, the current study focused primarily on cellular and signaling responses in vitro, and the translational significance of chronic alcohol-induced PARP inhibitor sensitivity will require further investigation in preclinical models and clinical specimens.

In summary, our study demonstrates that chronic alcohol exposure induces persistent DNA damage response signaling and sensitizes breast epithelial cells to PARP inhibition. These findings provide experimental evidence linking chronic alcohol-associated genomic stress to therapeutic vulnerability in DNA repair pathways and suggest that persistent replication stress induced by alcohol exposure may represent a previously underappreciated determinant of PARP inhibitor responsiveness.

## Data Availability Statement

All data is contained within the manuscript.

## Author Contributions

MZ: Data curation and analysis, writing-review & editing; ZM: Data curation and analysis, writing-review & editing; AP: Data curation and analysis, writing-review & editing; XY: Conceptualization, funding acquisition, investigation, project administration, writing – drafting, editing & review. All authors have read and agreed to the published version of the manuscript.

## Acknowledgments

The authors extend their appreciation to the funding agency and support from colleagues in this department.

